# On variational solutions for whole brain serial-section histology using the computational anatomy random orbit model

**DOI:** 10.1101/271692

**Authors:** Brian C. Lee, Daniel J. Tward, Partha P. Mitra, Michael I. Miller

**Affiliations:** Center for Imaging Science, Johns Hopkins University, Baltimore, MD, USA; Cold Spring Harbor Laboratory, Cold Spring Harbor, NY, USA

## Abstract

This paper presents a variational framework for dense diffeomorphic atlas-mapping onto high-throughput histology stacks at the 20 *μ*m meso-scale. The observed sections are modelled as Gaussian random fields conditioned on a sequence of unknown section by section rigid motions and unknown diffeomorphic transformation of a three-dimensional atlas. To regularize over the high-dimensionality of our parameter space (which is a product space of the rigid motion dimensions and the diffeomorphism dimensions), the histology stacks are modelled as arising from a first order Sobolev space smoothness prior. We show that the joint maximum a-posteriori, penalized-likelihood estimator of our high dimensional parameter space emerges as a joint optimization interleaving rigid motion estimation for histology restacking and large deformation diffeomorphic metric mapping to atlas coordinates. We show that joint optimization in this parameter space solves the classical curvature non-identifiability of the histology stacking problem. The algorithms are demonstrated on a collection of whole-brain histological image stacks from the Mouse Brain Architecture Project.

**Author Summary:** New developments in neural tracing techniques have motivated the widespread use of histology as a modality for exploring the circuitry of the brain. Automated mapping of pre-labeled atlases onto modern large datasets of histological imagery is a critical step for elucidating the brain’s neural circuitry and shape. This task is challenging as histological sections are imaged independently and the reconstruction of the unsectioned volume is nontrivial. Typically, neuroanatomists use reference volumes of the same subject (e.g. MRI) to guide reconstruction. However, obtaining reference imagery is often non-standard, as in high-throughput animal models like mouse histology. Others have proposed using anatomical atlases as guides, but have not accounted for the intrinsic nonlinear shape difference from atlas to subject. Our method addresses these limitations by jointly optimizing reconstruction informed by an atlas simultaneously with the nonlinear change of coordinates that encapsulates anatomical variation. This accounts for intrinsic shape differences and enables rigorous, direct comparisons of atlas and subject coordinates. Using simulations, we demonstrate that our method recovers the reconstruction parameters more accurately than atlas-free models and innately produces accurate segmentations from simultaneous atlas mapping. We also demonstrate our method on the Mouse Brain Architecture dataset, successfully mapping and reconstructing over 500 brains.

## 1 INTRODUCTION

### 1.1 Mapping brain circuitry

Recent advances in brain imaging [1, 2], methods to label neurons [3], and computational methods have brought about a new era of neuroanatomical research, with a focus on comprehensively mapping brain circuits [4]. Mapping whole-brain circuitry is important for three distinct reasons: scientific understanding of how the brain works, mechanistic understanding of neurological and neuropsychiatric disorders, and as a comparison point for artificial neural networks used in machine learning [5, 6].

Circuit mapping is technique limited, and falls into three broad scales corresponding to distinct imaging modalities - indirect mapping at a macroscopic scale corresponding to MRI-based methods [7], and direct mapping at light (LM) and electron microscopic (EM) scales. For MRI and LM data, atlas mapping is an important step in the analysis. Several approaches exist for gathering LM data at the whole brain level [8–10]. For some of these approaches (two-photon serial block-face imaging, knife edge scanning microscopy and light sheet microscopy for cleared brains) two-dimensional (2D) optical sections are acquired in three-dimensional (3D) registry with each other, so that the only computational step required is 3D volumetric registration of the individual brain data set to a canonical atlas. However, for classical neurohistological approaches using tissue sectioning followed by histochemical processing, the 2D sections are gathered independently and each section can undergo an arbitrary rotation and translation compared to the block face. This may be considered a disadvantage of the classical neu-roanatomical workflow, however the physical sectioning method followed by conventional histochemical analysis has certain important advantages. This allows for the full spectrum of histochemical stains, acquisition of physical sections for downstream molecular analyses, and processing for larger brains (upto and including whole human brains). Therefore it is necessary to perform an intermediate 2D to 3D registration step, where the individually acquired 2D sections are mutually co-registered into a 3D volume.

This paper develops a joint stack reconstruction and atlas mapping procedure that simultaneously restacks the 2D histology sections, applying a sequence of rigid motions to the sections, and estimates the diffeomorphic correspondence between the registered histology stack and the 3D atlas. We apply these algorithms to data sets from the Mouse Brain Architecture Project (MBAP), for which the experimental workflow generating the data utilizes a tape transfer technique [11], allowing for the sections to maintain geometrical rigidity within section and also allowing for physically disjoint components to maintain their spatial relations. The tape method ensures that the number of missing sections is minimal, with serial sections cut at a thickness of 20 *μ*m and alternate sections subjected to Nissl staining alongside staining with histochemical or fluorescent label. These Nissl stained sections form the basis of alignment to a Nissl whole-brain reference atlas.

### 1.2 Computational anatomy methods for brain histology

The histological reconstruction problem has been explored by several groups previously. Malandain first described the ill-posedness of reconstructing 3D sections and object curvature without prior knowledge of the shape of the object [12]. Rigid transformations for stack reconstruction have been estimated via block-matching of histological sections in [13], with point information based on landmarks introduced to guide volume reconstruction [14]. Dense external reference information such as MRI has been applied to guide reconstruction via registration of corresponding block-face photographs and for histology to MRI mapping [15, 16]. Anatomical atlases have also been suggested as guides for reconstruction, but without accounting for the intrinsic nonlinear shape differences between an atlas and any given subject brain [17].

The principal contribution of this work is to rigorously solve the problem when an external resource of identical geometry (such as an MRI of the same mouse) is not available, while accommodating for the innate anatomical variation from atlas to subject. The lack of a same-subject reference volume is often the standard in mouse brain histology and other large scale histology studies. This places us into the computational anatomy (CA) orbit problem for which constraints are inherited from an atlas that is diffeomorphic but not geometrically identical. With the availability of dense brain atlases at many resolution scales [18–21], methods to map atlas labels onto target coordinate systems are being ubiquitously deployed across neuroscience applications. Since Christensen’s early work [22], diffeomorphic transformation has become the de-facto standard as diffeomorphisms generate one-to-one and onto correspondences between coordinate systems. Herein we focus on the diffeomorphometry orbit model [23] of computational anatomy [24], where the space of dense volume imagery is modelled as a Riemannian orbit of an atlas under the diffeomorphism group. We use the large deformation diffeomorphic metric mapping (LDDMM) algorithm first derived for dense imagery by Beg [25] to retrieve the unknown high-dimensional reparameterization of the template coordinates.

Of course, for the histological stacking problem solved here, the interesting twist is the augmentation of the random orbit model with 3 rigid motion dimensions for each target section. At 20 *μ*m, this implies as many as 500 sections augmenting the high-dimensionality of the diffeomorphism space to include as many as 1500 extra dimensions for planar rigid motions for restacking. Here lies the crux of the challenge. To accommodate the high-dimensionality of the unknown rigid motions, the space of stacked targets is modelled to have finite-squared energy Sobolev norm, which enters the problem as a prior distribution restricting the roughness of the allowed restacked volumes. The variational method jointly optimizes over the highdimensional diffeomorphism associated to the atlas reparameterization and the high-dimensional concatenation of rigid motions associated to the target.

## 2 METHODS

### 2.1 The Log-Likelihood Model of the Histology Sectioning Problem

Figure 1 shows the components of the model for the histology stacking problem. We define the mouse brain to be sectioned as a dense three-dimensional (3D) object *I*(*x,y,z*), (*x,y,z*) ∈ ℝ^3^, modelled to be a smooth deformation of a known, given template *I*_0_ so that *I* = *I*_0_ ○ *φ*^−1^ for some invertible dif-feomorphic transformation *φ*. The Allen Institute’s mouse brain atlas [26] (CCF 2017) is taken as the template.

**Figure 1:**
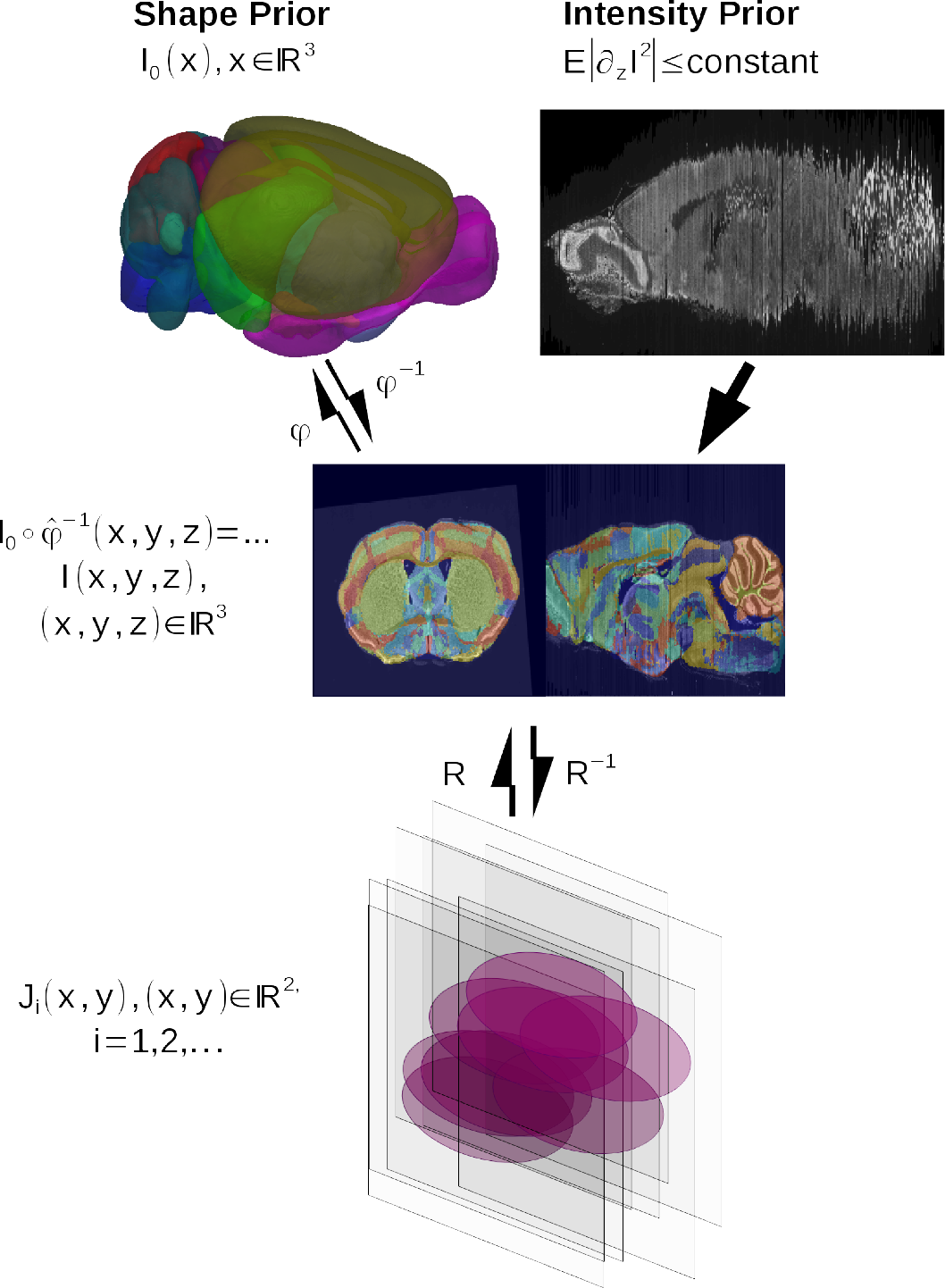
The histological sectioning model; the template *I*_0_, the mouse brain in the orbit 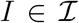 and observed histological sections *j*_*i*_, *i* = 1,…, *n*. The Sobolev image intensity prior and the shape prior are depicted in the top row. The model shows the template and mouse brain as elements of the same orbit *I*_0_, 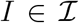, so there exists diffeomorphism *I* = *I*_0_ ○ *φ*^−1^, *φ* ∈ *Diff.*

Distinct from volumetric imaging such as MRI which delivers a dense 3D metric of the brain, the histology procedure (bottom row, Figure 1) consisting of sectioning, staining, and imaging generates a jitter process which randomly translates and rotates the stack sections. Denote the rigid motions acting on the 2D sectioning planes *R*_*i*_: ℝ^2^ → ℝ^2^, 
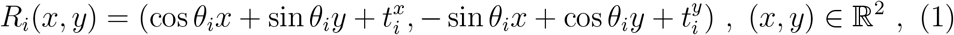
with *θ*_*i*_ the rotation angle and 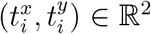 the translation vector in section *i*. The histology stack *J*_*i*_(*x,y*), (*x,y*) ∈ ℝ^2^,*i* = 1,…, *n*, is a sequence of 2D jittered image sections under smooth deformation of the atlas in noise: 
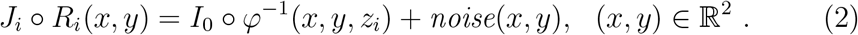

Modeling the photographic noise as Gaussian and conditioning on the sequences of jitters *R*_*i*_, *i* = 1,…, *n* and atlas deformation *I* = *I*_0_ ○ *φ*^−1^, *φ* ∈ *Diff*, the photographic sections *J*_*i*_ are a sequence of conditionally Gaussian random fields with log-likelihood 
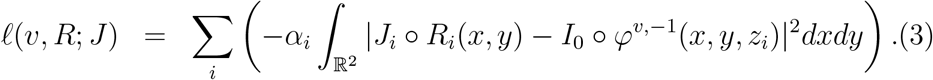

Here *α*_*i*_ is a weighting factor dependent on the noise of each section such that damaged sections can be weighted; *υ* denotes the vector field which indexes the deformation as a diffeomorphic flow (see below).

### 2.2 The Priors: Diffeomorphisms and Sobolev Smoothness of Images

The parameterization of the histology pipeline augments the standard random orbit model of computational anatomy with the rigid-motion dimensions of the random jitter sectioning process. The unknowns to be estimated become (*R*_1_,…, *R*_*n*_, *φ*) ∈ ℝ^3*n*^ × *Diff* for *n*—sections. At 20*μ*m then *n* = 500 implying the nuisance rigid motions are of high dimension *O*(1500). The solution space must be constrained. We use priors on the deformations and on the rigid motion stacking of the images.

#### The Diffeomorphism Prior

The histological stacking constrains the brains as smooth transformations of the template, where the diffeomorphisms are generated as diffeomorphic flows *φ*_*t*_ ∈ *Diff* [24], solving the ordinary differential equation

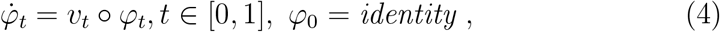
with *υ*_*t*_ the Eulerian velocity taking values in ℝ^3^, *identity* the identity mapping. The top row of Figure 1a shows that each *φ* has an inverse and that the random orbit model assumes any individual brain 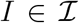 can be generated from the exemplar under the action of the diffeomorphism, so that for some *φ* ∈ *Diff*, *I* = *I* ○ *φ*^−1^. To score the distances between mouse brain coordinate systems and reject outlier solutions we use geodesic flows minimizing metric length [27]. Large deviations as measured by the diffeomorphometry metric [23] from template atlas to target mouse brain are thus removed from the solution space. The vector fields are modeled to be in a reproducing kernel Hilbert space (RKHS) (*V*, ‖ · ‖_*V*_), supporting one continuous spatial derivative, and having geodesic length between coordinate systems determined by the norm-square 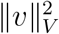 of the RKHS: 
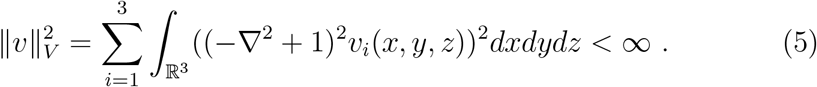

This square-metric is used as a quadratic potential for the smoothness prior between images *I*, 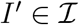 [28, 29] minimizes the action 
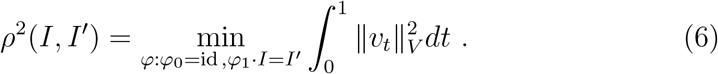

See Appendix B for the explicit equations for geodesics satisfying the Euler-Lagrange equations [27, 30] and Appendix A for the matrix Green’s kernel.

We use the notation *φ*^*υ*^ to emphasize the dependence of the diffeomor-phism and the geodesic metric on the vector field *υ*. Strictly speaking, the group generated by integrating (4) with finite norm ‖ · ‖_*V*_ is both dependent on the norm of *V* as well as a subgroup of all diffeomorphisms; we shall suppress that technical detail in the notation.

#### The Prior Distribution on Image Smoothness

To score the maximum a-posteriori (MAP) reconstruction of the rigid motions acting on the stack, we exploit a smoothness prior on the reconstructed histology stack which enforces the fact that anatomical structures are smooth and continuous. We model the images as arising from a smooth “Sobolev” or RKHS *I* ∈ *H*^*k*^ supporting derivatives 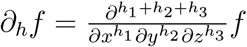 that are square integrable, with norm: 
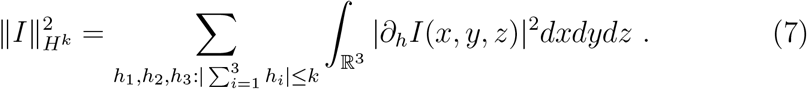

This is a quadratic form for a Gaussian random field prior on the dense histology stack with zero mean and covariance dependent on the squared norm 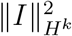. For the purpose of stacking, the z-axis sections are sparse 20 — 40*μ*m; the differential operators ∂_*h*_ are implemented via the difference operator along the sectioning z-axis (see Eqn. (8)). The Gaussian field has covariance determined by the difference operators; see [31] for example. We define the mixed differential-difference operator *D*_*h*_ as the centered difference for the z-partial derivatives, 
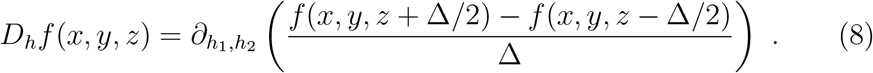

The gradient is forced to 0 at the boundaries of the image.

### 2.3 MAP, Penalized-Likelihood Reconstruction

Model the random sectioning with section-independent jitter as a product density 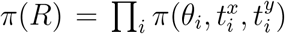, the priors centered at identity. Generating MAP estimates of the rigid motions generates the MAP estimator of the histology restacking problem denoted as 
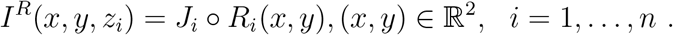

Since the diffeomorphisms are infinite dimensional, the maximization of the log-likelihood function with respect to a function with the deformation penalty is termed the “penalized-likelihood estimator”. Conditioned on the known atlas, the augmented random variables to be estimated are (*R*_1_,…, *R*_*n*_, *φ*) ∈ (ℝ^3*n*^ × *Diff*).

#### Problem 1 (MAP, Penalized-Likelihood Estimator).

*Given histology stack J*_*i*_(*x,y*), (*x,y*) ∈ ℝ^2^, *i* = 1,… *and reconstructed stack I*^*R*^(·, *z*_*i*_) = **J*_*i*_* ○ *R*_*i*_(·), *i* = 1,…, *n modelled as conditionally Gaussian random fields conditioned on jitter and smooth dormation of the template. The joint MAP, Penalized-Likelihood estimators* argmax_*R,υ*_ logπ(*R*, *υ*|*J*) *given by* 
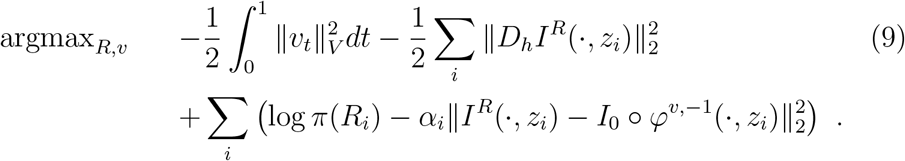

*The MAP, Penalized-Likelihood estimators satisfy* 
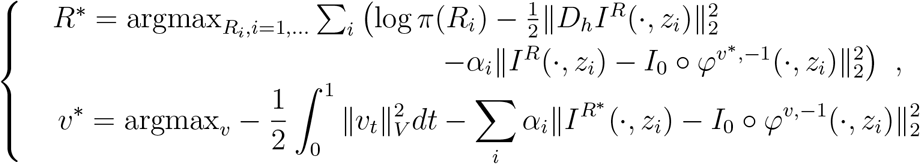
*with* 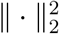 *denoting the norm per z-axis section:* 
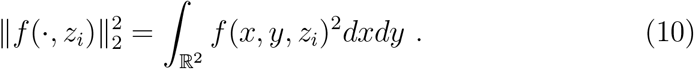

We call this the **atlas-informed** model. The first two prior terms of (9) control the smoothness of template deformation and the realigned target image stack, with the third keeping the rigid motions close to the identity. The last term is the “log-likelihood” conditioned on the other variables.

The optimization for the *R*\* rigid-motions is not decoupled across sections because of the smooth diffeomorphism of the LDDMM update and the Sobolev metric represented through the difference operator across the *z*-sections. The optimization of the vector field *υ*\* corresponds to the LDDMM solution of Beg [25].

The principal algorithm used for solving this joint MAP-penalized likelihood problem alternates between fixing the rigid motions and solving LD-DMM and fixing the diffeomorphism and solving for the rigid motions. This is described in the Methods section.

When there is no atlas available this is equivalent to setting *α_i_* small and becomes a MAP rigid motion restacking of the sections: 
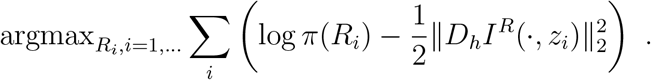

We term this the **atlas-free** model. The gradient of the rigid motions with respect to the components of translations *t*^*x*^,*t*^*y*^ and rotation *θ* is defined in Appendix C. The registration is not independent across sections due to coupling through the Sobolev metric.

### 2.4 Iterative Algorithm for Joint Penalized Likelihood and MAP Estimator

Here we describe the details of the algorithm used for solving for the MAP/penalized-likelihood problem of section 2.3. The algorithm alternately fixes the set of rigid motions while updating LDDMM and fixes the diffeomorphism while updating the rigid motions.

#### Algorithm 1.

0. *Initialize φ*^*new*^, *R*^*new*^ ← *φ^*init*^, *R*^*init*^, *I*^*old*^* ← *J* ○ *R*^*init*^:
1. 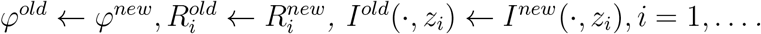
2. *Update LDDMM for diffeomorphic transformation of atlas coordinates:* 
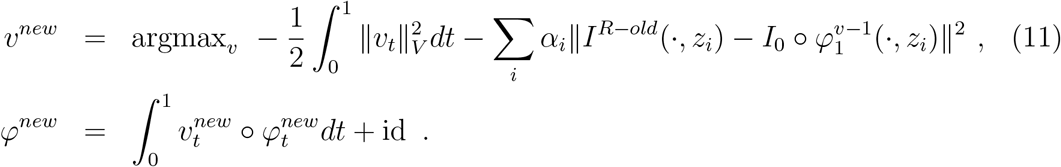
3. *Deform atlas I*_0_ ○ *φ*^*new*-1^ *and generate new histology image stack:* 
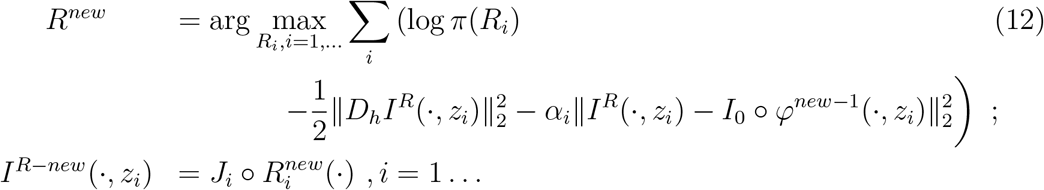
4. *Return to Step 1 until convergence criterion met.*

The form of the gradients for the rigid motions is given in Appendix D. The LDDMM update solutions are given by Beg [25].

### 2.5 Software Implementation

The algorithm described above is applied to Nissl histological stacks using the Allen Institute’s mouse brain atlas as a template. The Allen Mouse Brain Atlas is a micron-scale atlas that includes annotated Nissl-stained images at 10, 25, 50, and 100 *μ*m voxel resolution, with 738 labeled compartments in the annotation.

Atlas mapping is computed on the Nissl-stained histological image stack showing the clear definition of anatomical boundaries. The associated fluorescent tracer images are transformed to the Nissl stack so that the atlas subvolume labels can be cast onto the new modality. The fluorescent and Nissl images are registered within animals by applying rigid registration based on a mutual information cost function.

A software pipeline which performs start-to-finish registration operations was implemented on a high performance computing cluster for atlas-mapping and histology restacking on the Mouse Brain Architecture data. To date, the pipeline has been successfully run on over 1000 MBAP brains. The general pipeline workflow is illustrated in Figure 2.

**Figure 2:**
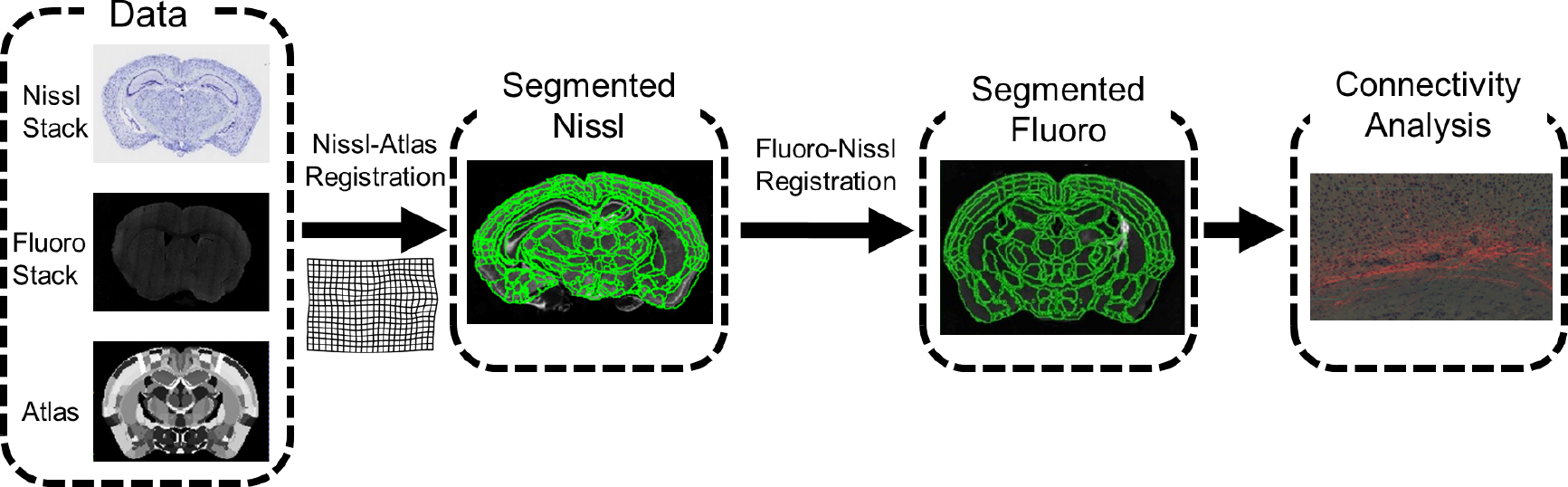
Histology registration pipeline workflow

The pipeline begins with a pre-selection of the atlas model. After affine registration of the Nissl sections is applied, the algorithm of section 2.4 is applied to compute the mapping between atlas and target coordinate spaces. In our application, we apply a two channel LDDMM [32] algorithm for the optimization with respect to *φ*, where the first channel is the Nissl-stained grayscale image, and the second channel is a mask of the brain tissue with ventricles and background set to a pixel value of zero. The brain mask for each brain stack is automatically generated by thresholding at an estimated background intensity value and applying morphological opening and closing for denoising. The threshold value is estimated by a RANSAC-like procedure over the image histogram, assuming a normal distribution of intensity values in the image foreground. A first-order Sobolev-norm (see below) is used for the smoothness constraint regularization of the histology stack. In order to accommodate for sections damaged by the histology process or structures excluded from imaging, the objective functions in all parts of the algorithm are optimized with respect to only the image data that exists. Essentially, this is a masking procedure on the cost function that allows matching between a whole atlas brain and some target which is a partial, or subset of a whole brain.

After registration of the structural Nissl image, the fluorescence volume is registered to its corresponding Nissl volume. The registration is restricted to rigid motions on each individual section. The optimization bears a similar form to equation (12) with the squared error matching term replaced with mutual information in order to account for the different modalities of the template and target histology stack. Once fluoro-to-Nissl registration is complete, the Nissl segmentation can be applied to the fluorescence image.

## 3 RESULTS AND DISCUSSION

### 3.1 Validation on Simulated Reconstructions

#### 3.1.1 The Phantom with Curvature

The model was applied to binary image phantoms in order to examine the “curvature” problem in which a 3D curved object cannot be accurately reconstructed after being sectioned. This is illustrated in Figure 3. We produced sections through the 3D phantom, applying the atlas-free and the atlas-informed models. The results from the atlas-free algorithm in which the sections are aligned based on the Sobolev smoothness followed by mapping of the atlas via LDDMM are summarized in Figure 3c. The atlas-free section alignment reconstructs the target stack, demonstrating a cylindrical reconstruction rather than the curved template shape, followed by LDDMM alignment *I*_0_ ○ *φ*^−1^. This illustrates the curvature issue. The atlas coordinate grid is transformed significantly (bottom right of Figure 3c) in order to match the target. Despite this significant deformation, there is some residual error in the atlas-to-target mapping with the remaining tendrils where the ends of the phantom did not shrink inwards. Here, the energy required to push the ends of the atlas inwards were greater than the potential image matching improvement.

Shown in Figure 3d is the atlas-informed solution. The bottom row of Figure 3d shows that simultaneously solving for reconstruction and registration parameters allows for more consistent stack reconstruction of the target resulting from the influence of the smooth deformation of the template onto the target in the joint solution.

**Figure 3:**
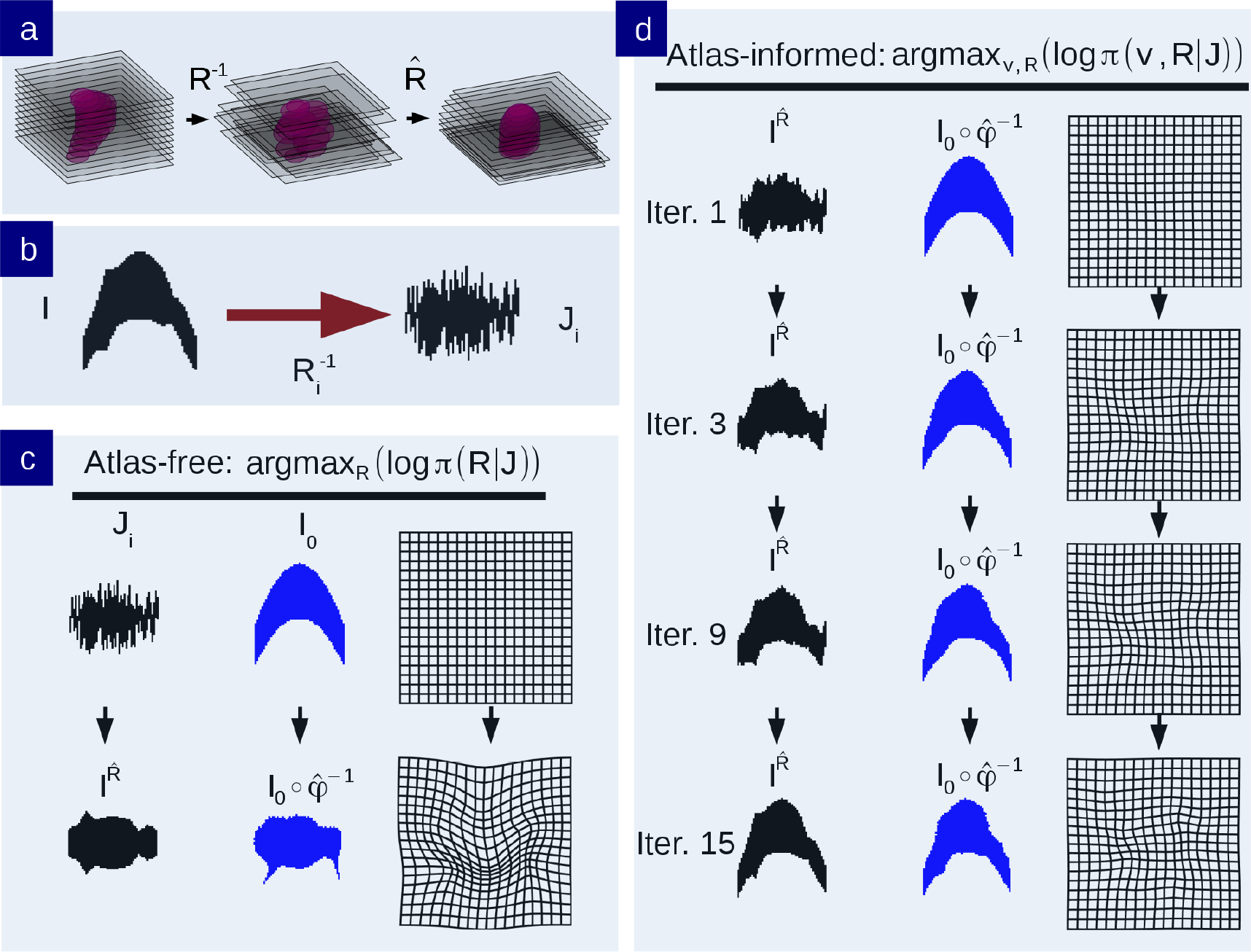
a) An illustration of the classic curvature reconstruction problem. b) The unobserved 3D-phantom is randomly sectioned and observed as *J*_*i,*_ *i* = 1,…, *n*. *c*) Reconstruction of the histological stack using the atlas-free method. The top row shows the histological stack and atlas. The bottom row shows the reconstructed histological stack 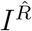 alongside the deformed phantom atlas *I* = *I*_0_ ○^−1^ which has been mapped to histological sections, and the diffeomorphic change of coordinates 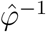. d) Reconstruction of phantom using the atlas-informed model. Each row depicts iterations of the reconstructed histological stack 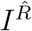 alongside the deformed atlas 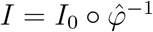 and deformed coordinates. The bottom row shows the convergence point of the algorithm.

These results are depicted by the difference in the motions of the atlas coordinate grids when deforming onto the targets in Figure 4. Tandem op-timization of section alignment parameters and diffeomorphisms produces a nonlinear mapping with lower metric cost (Fig 4c is less warped than Fig 4b).

**Figure 4:**
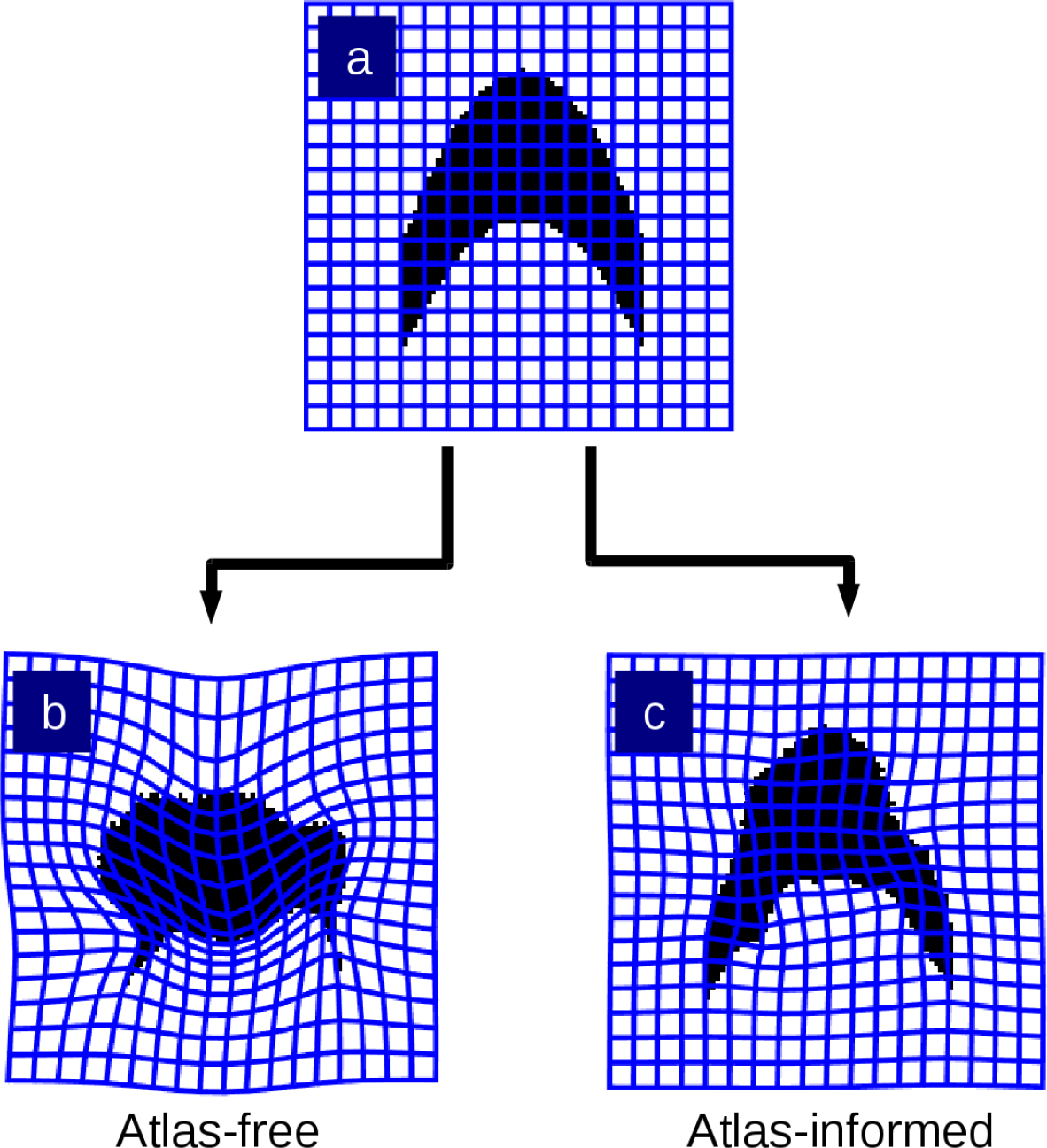
Transformed grids illustrating the difference in the mapping deformation from the atlas-free methods from (A) to histology stack target (B) versus the atlas-informed algorithm which produces (C).

#### 3.1.2 Jittering the Allen Atlas

A similar experiment was performed using the Allen mouse brain atlas as the 3D phantom. A target histology stack was generated by sectioning the Allen atlas in simulation and applying random rigid transforms to its coronal sections. The atlas images were sampled at 40 *μ*m isotropic voxels. This is depicted in Figure 5a. A simulated atlas was generated by applying a given random diffeomorphism to the Allen atlas. This random diffeomorphism is depicted in Figure 5c. The histology stacks were then reconstructed and diffeomorphic transformations generated between the atlas and target stacks using both models, intending to recover both the unknown rigid transforms from Figure 5a and the unknown diffeomorphism from Figure 5c. Figure 5b shows the atlas-free method method (bottom left) compared to the atlas-informed method (bottom right). The atlas-informed method nearly reproduces the original coordinates whereas the atlas-free method drifts away from the original coordinates. Note that although the diffeomorphisms are not identical, this does not necessarily indicate segmentation error as small differences in stack alignment can be compensated for by nonlinear registration during atlas-mapping.

**Figure 5:**
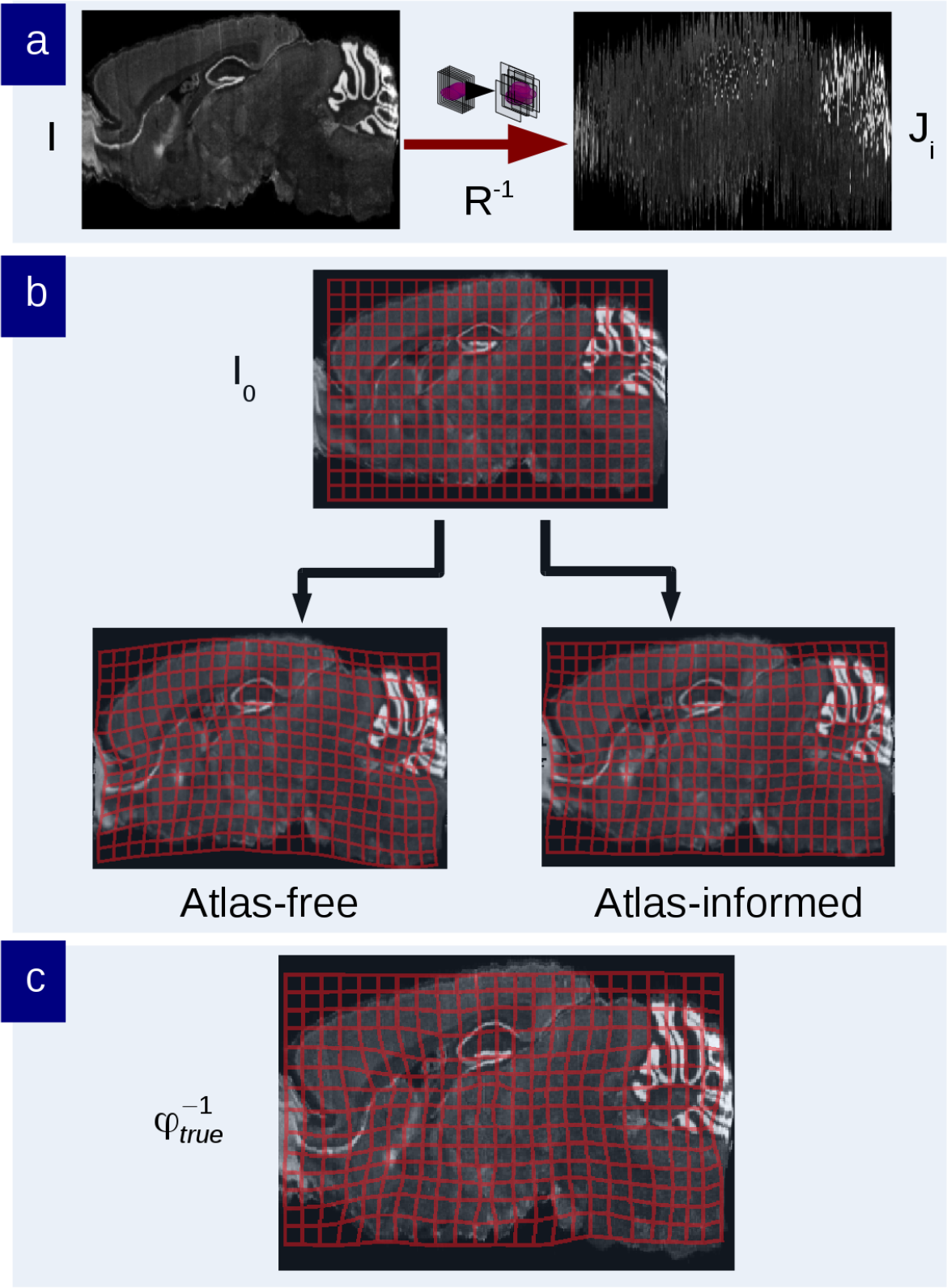
Atlas phantom simulation to validate recovery of sectioning parameters and diffeomorphic shape difference. a) The ground truth target *I* is sectioned to generate the observed target *J*_*i*_. b) Transformed grids illustrating the brain phantom atlas (top) shown mapped onto the histological stack using the atlas-free algorithm (bottom left) and the atlas-informed algorithm (bottom right). c) The ground truth diffeomorphism to be recovered.

#### 3.1.3 Simulated Bias and Variance Statistics

Figure 6 and 7 show results quantifying the bias and viarance of the joint estimation of the diffeomorphism transformation and the rigid motion jitter in simulation. Eqn. (2) was simulated over a range of Gaussian white noise selections while simultaneously varying the jitter rigid motions of the sections along with multiple deformations of shearing applied to the template *I*_0_. Shearing produced images where each section was successively offset by 0.25 pixels in both x and y directions, cumulatively producing the “shear” effect illustrated in Figure 6. Figure 7b keeps the stack jitter fixed and varies the noise levels; Figure 7c varies the stack jitter. The RMSE, bias, and standard deviation of the estimated parameters were computed in each experiment and plotted as a function of error units versus noise level. 500 simulations per experiment were performed.

**Figure 6:**
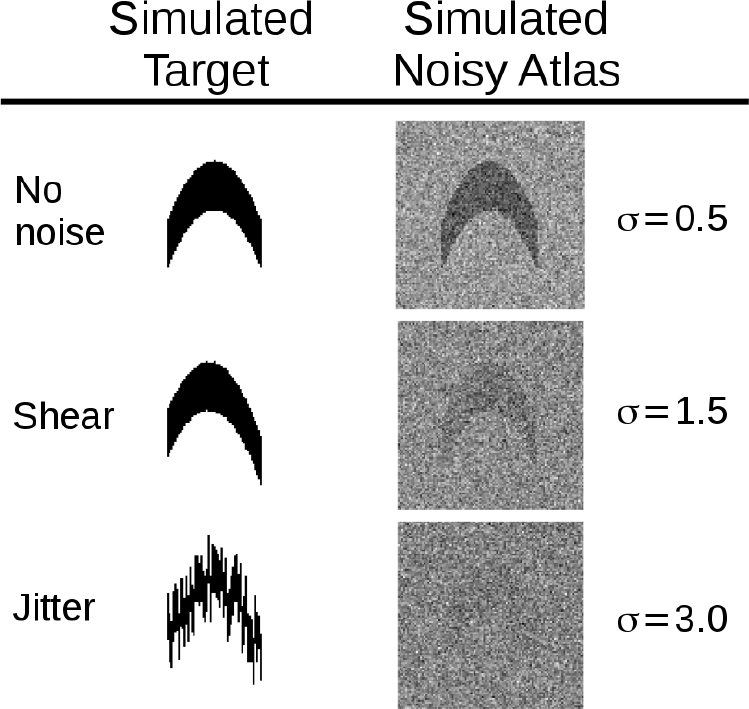
Left column shows phantom for identity, shearing, and jitter of sections (successive rows); right column shows Gaussian white noise added to the atlas at various standard deviations. The jitter random rigid motions were normally distributed 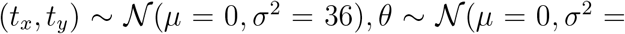 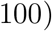 in pixel units.

**Figure 7:**
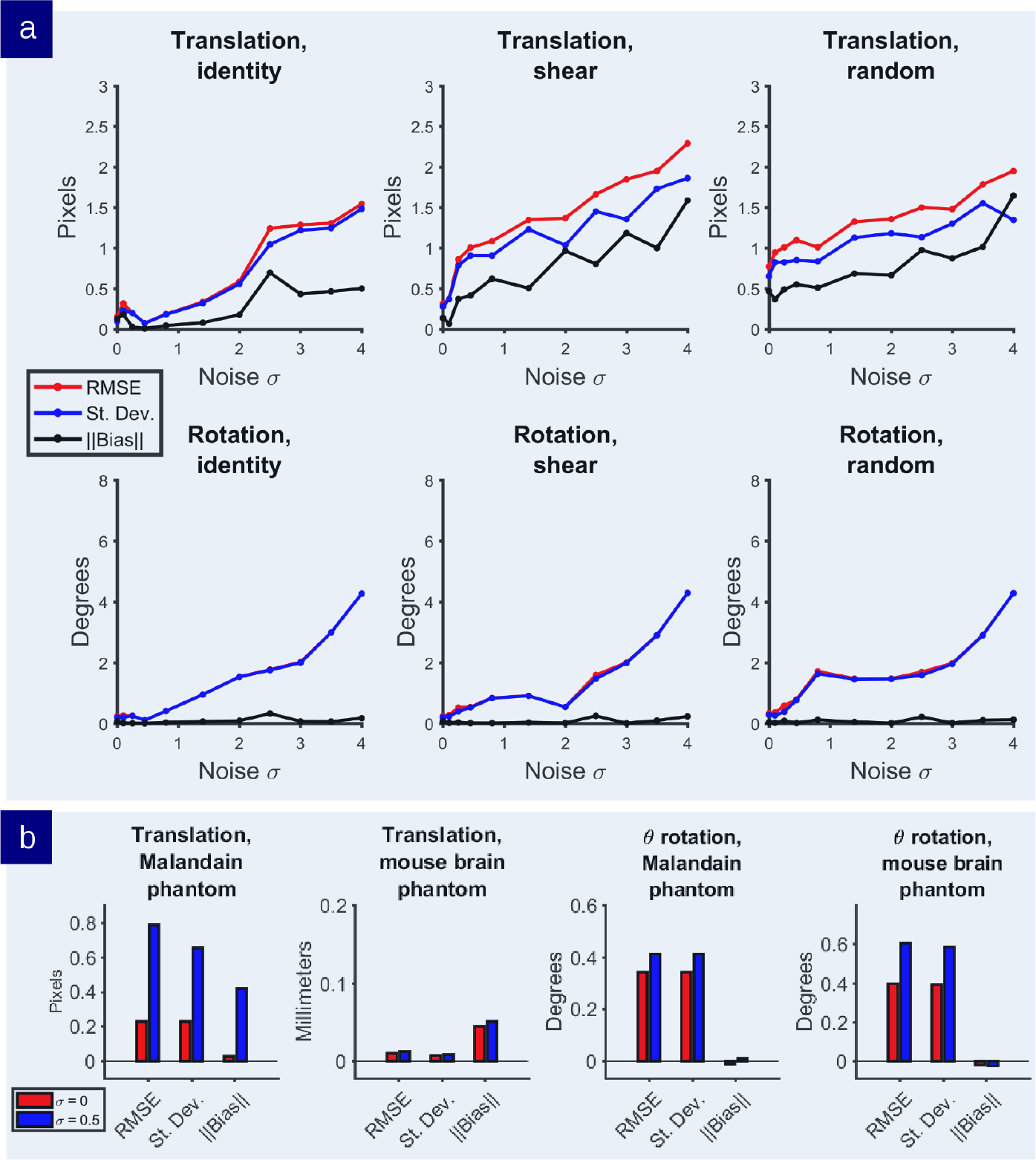
a) Statistics on the translation-rotation estimators for noise levels varying initial conditions. b) Statistics on the rigid motion estimators where the section jitter was added in a random fashion.

In each experiment, estimator accuracy is preserved up to high noise levels. At typical noise levels (*σ* ≤ 0.5), we observe subpixel RMSE and small bias. Figure 7b shows that the rotation estimator is virtually unbiased whereas the translation estimator does have small subvoxel bias. It is likely that more rotational error is accounted for by section realignment than deformable mapping, whereas both play a relatively balanced role in translation correction. Small motions are ill-posed in that small rigid-motions can accommodate small atlas deformation. Figure 7c (top row) shows the case where the target stack are jittered. Estimator statistics are computed in each of these cases showing similar subpixel errors.

Simulations examining bias and variance were also run on the Allen atlas brain as the phantom. The reconstruction RMSE observed in the brain phantom simulation (bottom row of Figure 7c) is lower than that observed in the simple curved phantom in pixels. It is likely that this is due to the presence of more contour lines in grayscale images versus binary images. These additional features allow for more accurate distinction of matching error than simpler images with small numbers of distinct level lnes. This is consistent with the demonstration in [27] showing that the stabilizer of the group corresponding to vector fields tangent to the level lines of the image cannot be uniquely identified or retrieved via any mapping methods that look at color or contrast of the image as the identifying feature.

### 3.2 Mouse Brain Architecture Project data

A final experiment was conducted on brain data sampled from the MBAP database, using the Allen mouse brain as the atlas. We selected specific targets which were prone to poor registration results due to image intensity local minima. In particular, structures like the cerebellum tend to be difficult to register accurately due to their folded nature; one fold can easily be mistaken for the adjacent fold, and if the target and atlas are not well initialized, the deformation required to flow one fold onto another can have a high metric cost. We are also interested in inspecting lower-contrast structures like the corpus callossum, which may be poorly registered due to local minima in other nearby bright structures.

**Figure 8:**
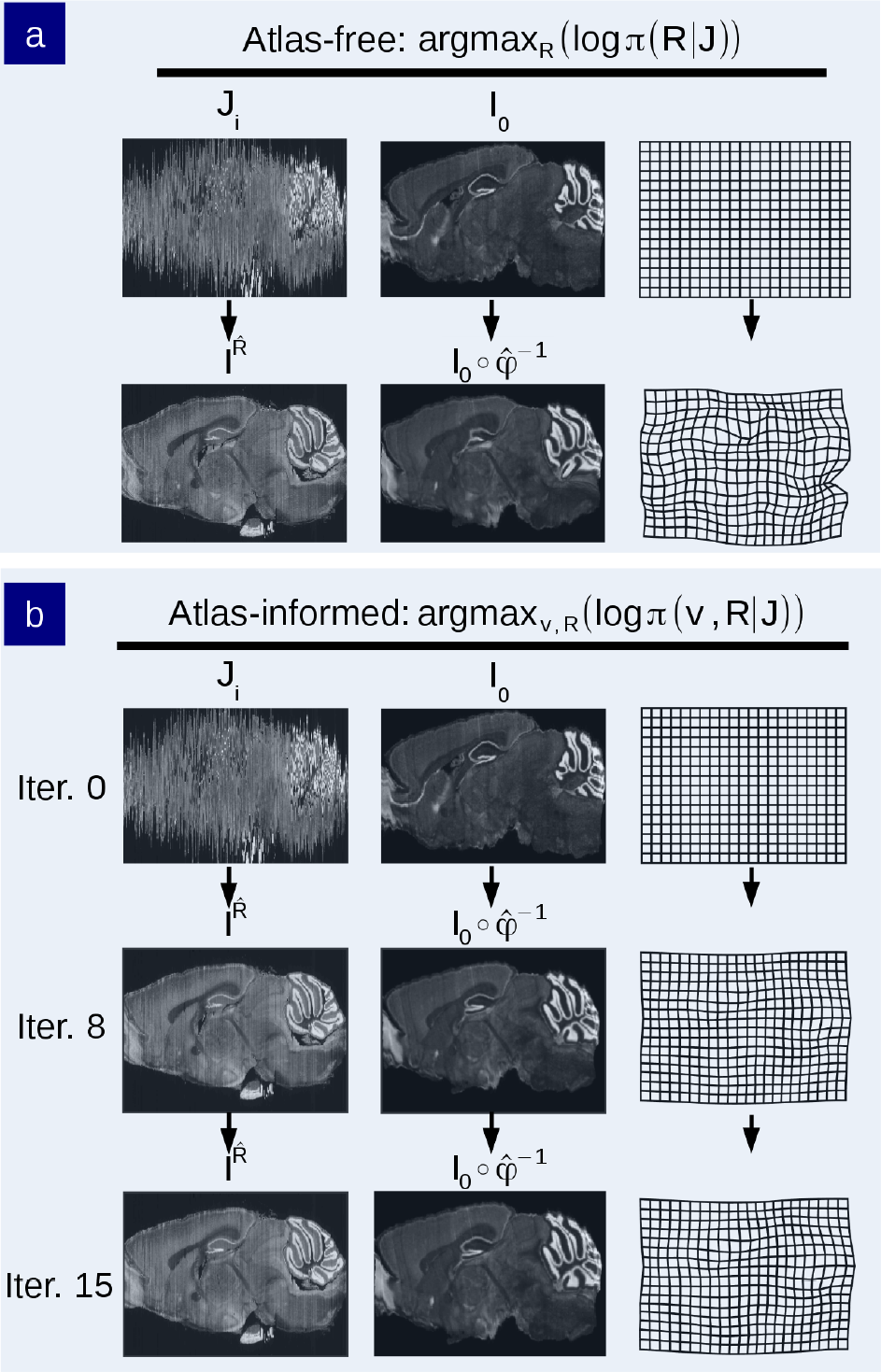
a) Reconstruction of an MBA Nissl-stained brain histological stack using the atlas-free method. Top row shows the histological stack and Allen mouse brain atlas. Bottom row shows the reconstructed histological stack 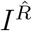 alongside the deformed phantom atlas *I*, and the diffeomorphic change of coordinates (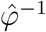. b) Reconstruction using the atlas-free method. Top row shows the histological stack and Allen mouse brain atlas. Middle row depicts intermediate iterations of the reconstructed stack 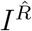 alongside the deformed atlas 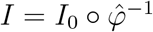 and coordinate grid. Bottom row shows the convergence point of algorithm.

The reconstructed histological target stack in the atlas-informed model shown in Figure 8a takes on the shape of the atlas but is prone to reconstruction artifacts. The deformation grids produced by the atlas-informed mapping is much smoother and has many fewer wrinkles than the atlas-free mapping. This is seen clearly in Figure 9.

**Figure 9:**
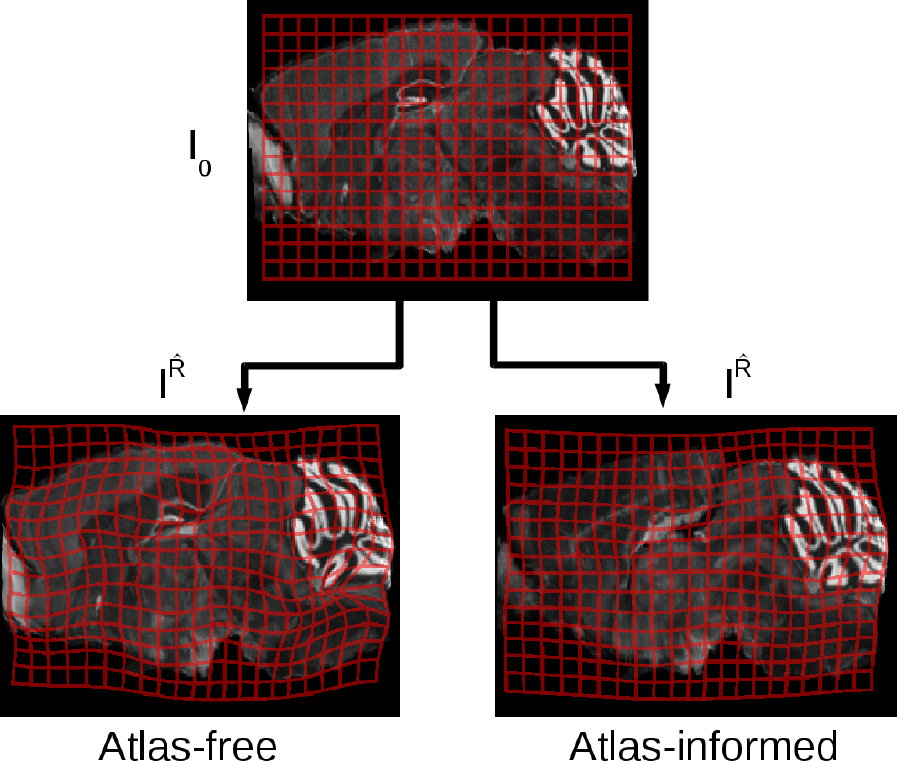
Transformed grids illustrating the difference in the mapping deformation from atlas (top) to target using the atlas-free method (bottom left) versus the atlas-informed method (bottom right), performed on real brain data from the MBA Project.

Figure 10 shows examples of improved segmentations in selected regions of the brain. The atlas-informed model generates more accurate segmentation results and produces smoother mappings as exhibited by the less wrinkled and distorted grids (bottom row b), showing more consistent results throughout the MBAP dataset.

**Figure 10:**
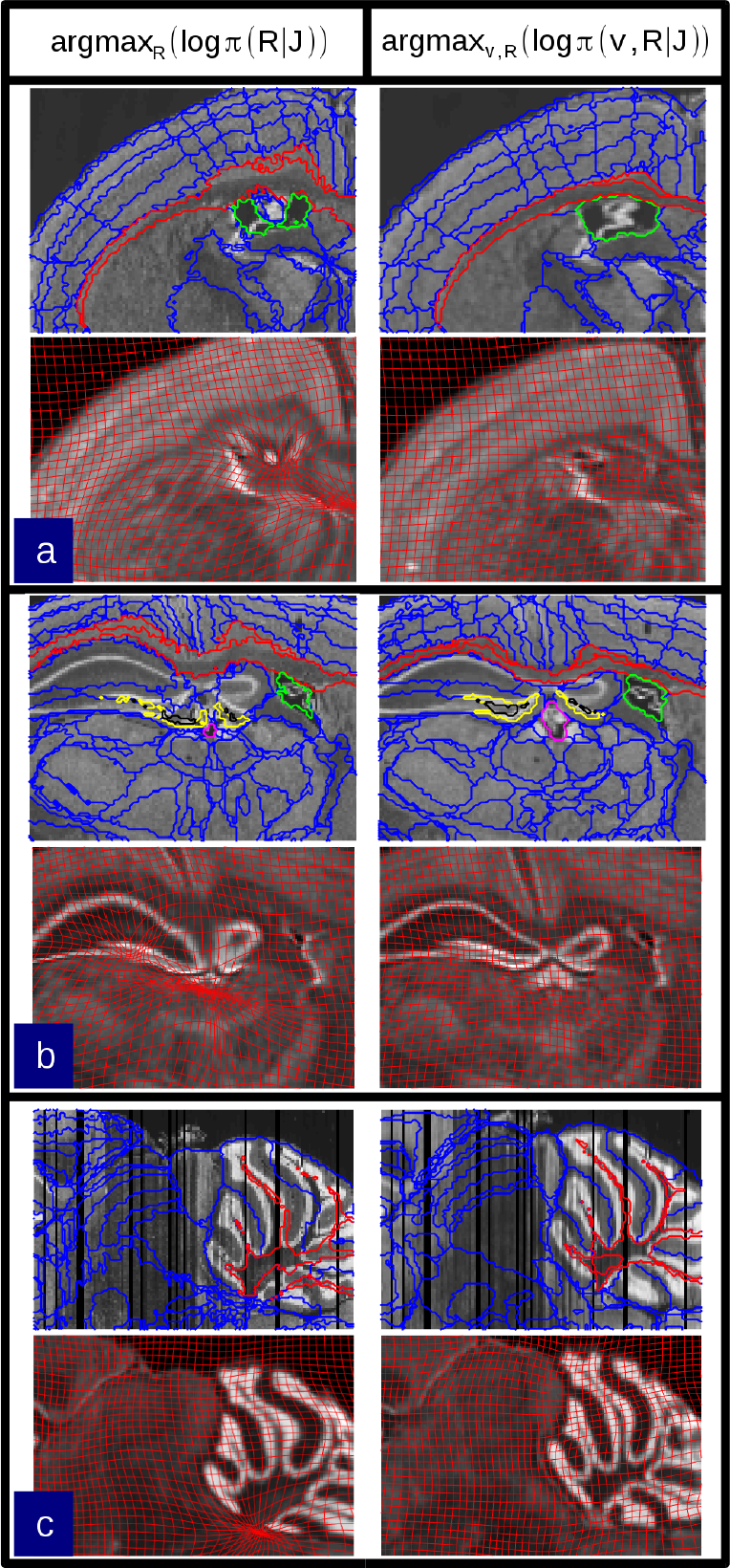
Selected regions of the brain segmented by the atlas-informed and atlas-free models carry the label map from the Allen atlas. The left column shows several examples where optimization of the atlas-free solution is trapped in false minima due to folded or low-contrast structures. The right column shows correction by the atlas-informed algorithm. A) The corpus callossum and lateral ventricle. B) The dentate gyrus, corpus callossum, and lateral ventricle. C) The cerebellar white matter.

## 4 CONCLUSION

This paper examines the CA random orbit model at the mesoscale for the stacking of sectioned whole brains coupled with mapping to annotated atlases. The standard CA model has been expanded to include the O(3 × *n*) extra rigid motion dimensions representing the planar histology sections. The estimation procedure solved here simultaneously estimates the diffeomorphic change of coordinates between atlas and target histological stack, as well as the “nuisance” rigid motion parameters for each section in stack space. This requires the introduction of a smoothness constraint on the target jitter simultaneous with LDDMM, which is enforced via a Sobolev metric, encouraging the reconstructed stack to be smooth by controlling the derivative along the cutting axis.

Results are shown demonstrating that the introduction of an atlas into the estimation scheme solves several of the classic problems associated with volume reconstruction, including the reconstructin of the curvature of extended structures. Since the atlas gives *a priori* indication of the global shape, the tendency to remove distortions along the section axis is balanced against the desire to minimize the amount of deformation of the atlas onto the reconstruction. The algorithm is shown to mediate this tension well.

## Acknowledgments

The authors would like to acknowledge Keerthi K Ram and Daniel Fer-rante for all of their guidance in understanding the Mouse Brain Architecture Project datasets. This work was supported by the G. Harold and Leila Y. Mathers Foundation, the Crick-Clay Fellowship, the H.N. Mahabala Chair, National Science Foundation Eager award 1450957, the Computational Anatomy Science Gateway as part of the Extreme Science and Engineering Discovery Environment (XSEDE) with grant number ASC140026, NIH DA036400, as well as the Kavli Neuroscience Discovery Institute supported by the Kavli Foundation.

## Supporting Information

### A Reproducing Kernel Hilbert Space and Green’s Kernel

The Green’s kernel is translation invariant and takes the form

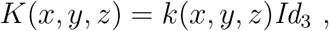
with *Id*_3_ the 3 × 3 identity matrix, for the Green’s function continuously differentiable: 
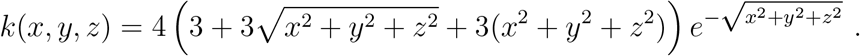

This Green’s function satisfies (−∇^2^ + 1)^3^*k*(*x, y, z*) = δ(*x, y, z*), where (−∇^2^ + 1)^4^ is referred to as *A*. The reproducing kernel Hilbert space (RKHS) with this Green’s kernel corresponds to vector fields satisfying 
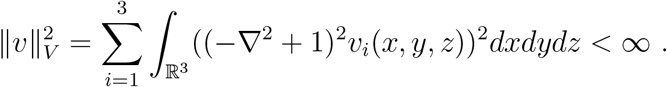

### B Geodesics solving Euler-Lagrange Equations

The explicit equations for geodesics associated to the RKHS norm ‖*υ*‖_*V*_ and the geodesics satisfy the Euler-Lagrange equations [27, 30] given by the triple of equations. 
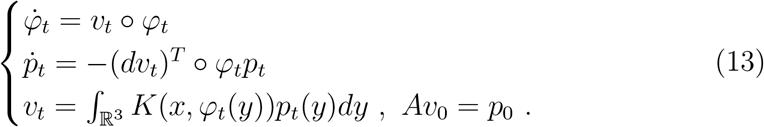

To prove the Hamiltonian momentum evolution the second equation *ṗ* = – (*dυ*)^*T*^ ○ *φp* of (13) for *Aυ* a classical function we use the inner product notation ⟨ ·, · ⟩ to calculate the Lagrangian: 
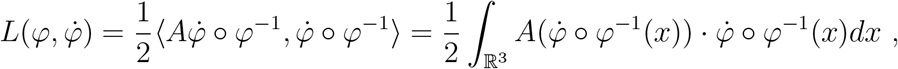
with the variation giving the Euler-Lagrange equations: 
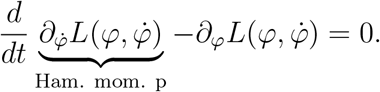

To get the Hamiltonian momentum 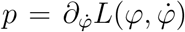, we take variation with respect to Lagrangian velocity 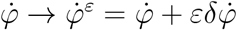 and *φ* → *φ* + *ɛ δ φ* giving 
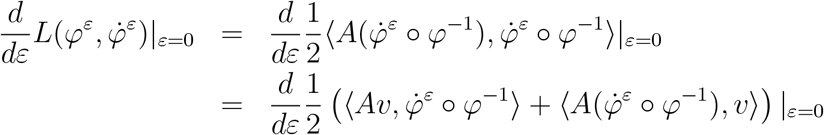

Combining gives the Hamiltonian momentum : 
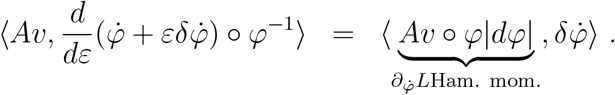

The variation *φ* → *φ*^*ɛ*^ = *φ* + *ɛ δ φ* requires the inverse: 
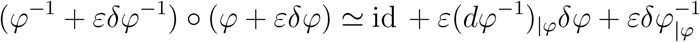
which gives first order perturbation 
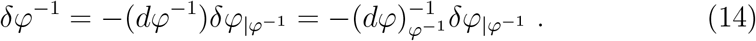

Taking a similar variation of the Lagrangian as above but with respect to the Lagrangian velocity gives 
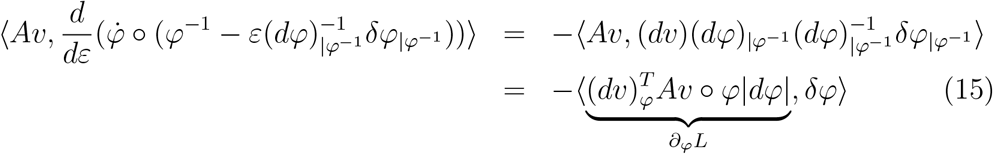

The third equation of (13) follows from *p* = *Aυ* ○ *φ*|*dφ*|. Integrating with the Green’s kernel gives the expression *υ*_*t*_(·) = ∫ *K*(·, *φ*_*t*_(*y*))*p*_*t*_(*y*)*dy*.

### C Gradients for Atlas Free Model

We can write the gradient with respect to the components of *R* (translation vector *t* and rotation matrix *r* parametrized by rotation angle *θ* and section number *z*), where ∇_*X*_ is the 2D in-plane gradient, *σ*_*JJ*_ is the weighting factor on the image smoothness prior. Rotations and translations are penalized by a regularization prior centered at identity (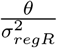 and 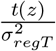, respectively), where *σ*_*regR*_ and *σ*_*regT*_ are weighting factors on the rotation and translation priors.

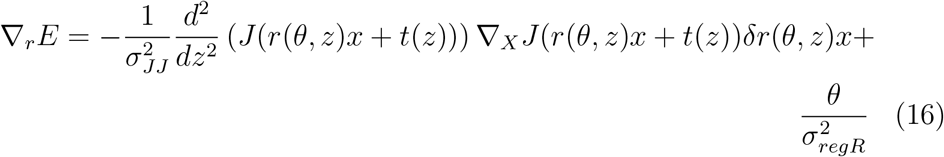

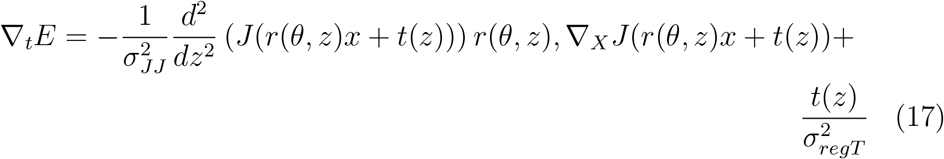

### D Gradients for Random Orbit Model

The minimization of the energy *E*_*υ*_ of (11) in terms of the vector field is the LDDMM gradient of Beg [25]: 
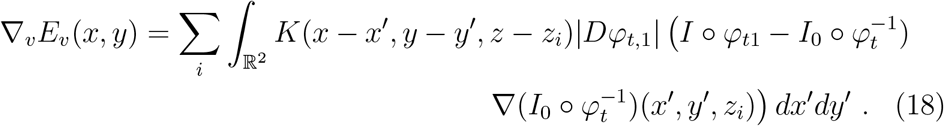

#### D.1 Variation of the Image Matching Term

The variation of ∫ (*I* – *I*_0_ ○ *φ*^−1^)^2^*dx* via perturbation *φ* → ^*ɛ*^ = *φ* + *ɛ δ φ* requires the inverse perturbation 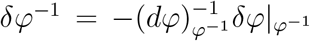, derived in (14) above. Then we have 
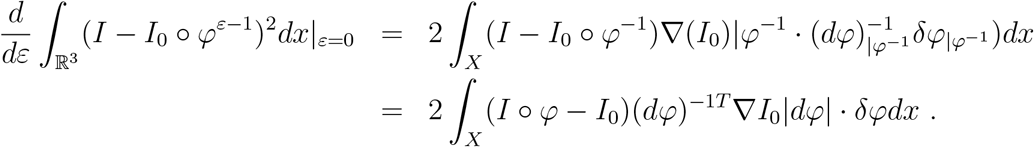

#### D.2 Rigid motion variations

Rigid motion minimization is standard for rigid registration in 2D and 3D images. Denoting 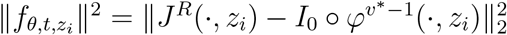 to represent each rigid registration norm-square minimization within each histological plane, then 
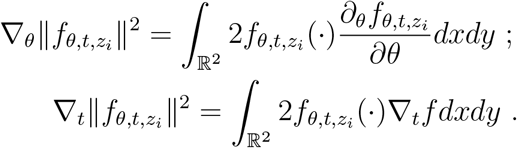

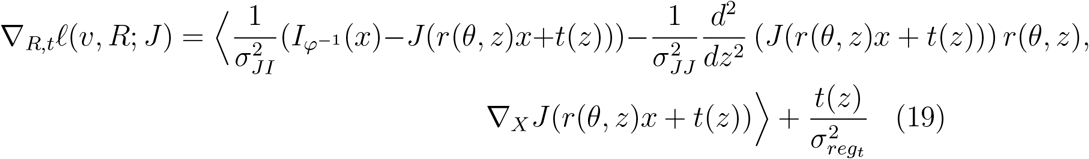

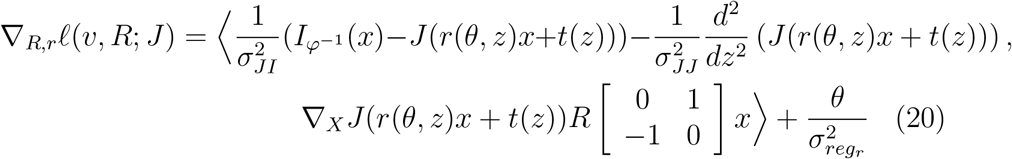
where *σ*_*JI*_ is a weighting factor on the matching term between atlas and target.

## References

[1] Hagmann P, Cammoun L, Gigandet X, Meuli R, Honey CJ, Wedeen VJ, et al. Mapping the structural core of human cerebral cortex. PLoS Biology. 2008;6(7). doi:10.1371/journal.pbio.0060159.

[2] Rubinov M, Sporns O. Complex network measures of brain connectivity: uses and interpretations. Neuroimage. 2010;52(3):1059–1069. doi:10.1016/j.neuroimage.2009.10.003.

[3] Taniguchi H, He M, Wu P, Kim S, Paik R, Sugino K, et al. A resource of cre driver lines for genetic targeting of gabaergic neurons in cerebral cortex. Neuron. 2011;71(6):995–1013. doi:10.1016/j.neuron.2011.07.026.

[4] Kasthuri N, Lichtman JW. The rise of the ‘projectome’. Nature Methods. 2007;4(4):307–309. doi:10.1038/nmeth0407-307.

[5] Mitra PP. The circuit architecture of whole brains at the mesoscopic scale. Neuron. 2014;83(6):1273–1283. doi:10.1016/j.neuron.2014.08.055.

[6] Sporns O. The human connectome: a complex network. Annals of the New York Academy of Sciences. 2011;1224(1):109–125. doi:10.1111/j.1749-6632.2010.05888.x.

[7] Hagmann P, Kurant M, Gigandet X, Thiran P, Wedeen VJ, Meuli R, et al. Mapping human whole-brain structural networks with diffusion MRI. PLoS One. 2007;2(7). doi:10.1371/journal.pone.0000597.

[8] Osten P, Margrie TW. Mapping brain circuitry with a light microscope. Nature Methods. 2013;10(6):515–523. doi:10.1038/nmeth.2477.

[9] Okano H, Mitra PP. Brain-mapping projects using the common marmoset. Neuroscience research. 2015;93:3–7. doi:10.1016/j.neures.2014.08.014.

[10] Papp EA, Leergaard TB, Csucs G, Bjaalie JG. Brain-wide mapping of axonal connections: workflow for automated detection and spatial analysis of labeling in microscopic sections. Frontiers in neuroinformatics. 2016;10. doi:10.3389/fninf.2016.00011.

[11] Pinksly V, Jones J, Tolpygo AS, Franciotti N, Weber K, Mitra PP. High-Throughput Method of Whole-Brain Sectioning, Using the Tape-Transfer Technique. PLoS One. 2015;10(7). doi:10.1371/journal.pone.0102363.

[12] Malandain G, Bardinet E, Nelissen K, Vanduffel W. Fusion of autoradiographs with an MR volume using 2-D and 3-D linear transformations. Neuroimage. 2004;23. doi:10.1016/j.neuroimage.2004.04.038.

[13] Ourselin S, Roche A, Subsol G, Pennec X, Ayache N. Reconstructing a 3D structure from serial histological sections. Image Vision Comput. Image Vision Comput. 2001;19(1). doi:10.1016/S0262-8856(00)00052-4.

[14] Streicher J, Weninger W, Muller G. External marker-based automatic congruencing: a new method of 3D reconstruction from serial sections. Anat Rec. 1997;248(4):583–602. doi:10.1002/(SICI)1097-0185(199708)248:4j583::AID-AR10¿3.0.C0;2-L.

[15] Dauguet J, Delzescaux T, Conde F, Mangin J, Ayache N, Hantraye P. Three-dimensional reconstruction of stained histological slices and 3D non-linear registration with in vivo MRI for whole baboon brain. J Neurosci Methods. 2007;164(1):191–204. doi:10.1016/j.jneumeth.2007.04.017.

[16] Adler DH, Pluta J, Kadivar S, Craige C, Gee JC, Avants BB, et al. Histology-derived volumetric annotation of the human hippocampal subfields in postmortem MRI. Neuroimage. 2014;84:505–23. doi:10.1016/j.neuroimage.2013.08.067.

[17] Qiu X, Shin L, Pridmore T, Pitiot A, Wang D. Atlas-guided correction of brain histology distortion. J Pathol Inform. 2011;2(S7). doi:10.4103/2153-3539.92038.

[18] Mai JK, Paxinos G. The Human Nervous System. Academic Press; 2011.

[19] Mori S, Oishi K, Faria A, Zijl PCM. MRI Atlas of Human White Matter. Elsevier; 2005.

[20] Chuang N, Mori S, Yamamoto A, Jiang H, Ye X, Xu X, et al. An MRI-based atlas and database of the developing mouse brain. Neuroimage. 2011;54(1):80–89. doi:10.1016/j.neuroimage.2010.07.043.

[21] Mori S, Oishi K, Faria AV, Miller MI. Atlas-Based Neuroinformatics via MRI: Harnessing Information from Past Clinical Cases and Quantitative Image Analysis for Patient Care. Annual Review of Biomedical Engineering. 2013;15:71–92. doi:10.1146/annurev-bioeng-071812-152335.

[22] Christensen G, Miller MI, Rabbit RD. Deformable templates using large deformation kinematics. IEEE Transactions of Medical Imaging. 1995;5:1435–1447. doi:10.1109/83.536892.

[23] Miller MI, Trouvé A, Younes L. Diffeomorphometry and geodesic positioning systems for human anatomy. Technology. 2014;1:36.

[24] Grenander U, Miller MI. Computational anatomy: An emerging discipline. Quarterly of Applied Mathematics. 1998;56(4):617–694. doi:10.1090/qam/1668732.

[25] Beg MF, Miller MI, Trouve A, Younes L. Computing Large Deformation Metric Mappings via Geodesic Flows of Diffeomor-phisms. International Journal of Computer Vision. 2005;61(2):139–157. doi:10.1023/B:VISI.0000043755.93987.aa.

[26] Lein ES, et al. Genome-wide atlas of gene expression in the adult mouse brain. Nature. 2007;445:168–176. doi:10.1038/nature05453.

[27] Miller MI, Trouvé A, Younes L. Geodesic Shooting for Computational Anatomy. Journal of Mathematical Imaging and Vision. 2006;24(2):209–228. doi:10.1007/s10851-005-3624-0.

[28] Miller M, Younes L. Group Actions, Homeomorphisms, and Matching: A General Framework. International Journal of Computer Vision. 2001;41(4):61–84. doi:10.1023/A:1011161132514.

[29] Miller MI, Trouvé A, Younes L. Hamiltonian Systems and Optimal Control in Computational Anatomy: 100 Years Since D’Arcy Thompson. Annual Review of Biomed Engineering. 2015;(17):447–509. doi:10.1146/annurev-bioeng-071114-040601.

[30] Miller MI, Trouve A, Younes L. On the metrics and euler-lagrange equations of computational anatomy. Annual Review of Biomedical Engineering. 2002;4(1):375–405. doi:10.1146/annurev.bioeng.4.092101.125733.

[31] Lanterman AD, Grenander U, Miller MI. Bayesian Segmentation via Asymptotic Partition Functions. IEEE Trans on Pattern Analysis and Machine Intelligence. 2000;22(4):337–347. doi:10.1109/34.845376.

[32] Ceritoglu C, Oishi K, Li X, Chou M, Younes L, Albert M, et al. Multi-contrast large deformation diffeomorphic metric mapping for diffusion tensor imaging. Neuroimage. 2009;47(2):618–27. doi:10.1016/j.neuroimage.2009.04.057.

